# Uncovering genes involved in pollinator-driven mating system shifts and selfing syndrome evolution in *Brassica rapa*

**DOI:** 10.1101/2023.09.29.560147

**Authors:** Xeniya V. Kofler, Ueli Grossniklaus, Florian P. Schiestl, Léa Frachon

## Abstract

**Background:** Shifts in pollinator occurrence and their pollen transport effectiveness drive the evolution of mating systems in flowering plants. A decline in pollinator numbers can lead to the phenotypic evolution of floral traits favoring self-pollination (selfing syndrome). Understanding the genomic basis involved in such shifts of the mating system is crucial for predicting a species’ persistence or extinction under changing biotic and abiotic stressors in natural populations. We investigated loci showing high genetic differentiation before and after selection in fast-cycling *Brassica rapa* that were associated with rapid phenotypic evolution toward the selfing syndrome (reduction in petal width, stamen and pistil length, and herkogamy). Combining a genotyping-by-sequencing (GBS) approach with a genome-wide association study (GWAS), we shed light onto the genetic basis associated with the mating system shift over nine generations of pollination by *Episyphrus balteatus* (hoverflies), an abundant pollinator of *B. rapa*. Moreover, we functionally validated the involvement of candidate genes associated with changes in floral morphology by studying corresponding mutants in the model plant *Arabidopsis thaliana*.

**Results:** We found that the mating system of *B. rapa* shifted from predominantly outcrossing to mixed mating with high rates of autonomous selfing, accompanied by a rapid reduction of floral morphological traits and herkogamy involving many loci. We found 31 candidate genes associated with the affected traits that were involved in a wide range of functions from DNA/RNA binding to transport. Our functional validation in *A. thaliana* confirmed that four of the identified genes are indeed involved in regulating the size of floral organs. Interestingly, two genes, both coding for the same protein complex responsible for active DNA methylation were successfully validated and closely linked to two correlated phenotypic traits, namely pistil and stamen length.

**Conclusions:** Altogether, our study shows that hoverfly pollination leads to rapid evolutionary changes of the mating system through polygenic changes, highlighting the importance of using genomic approaches to understand pollinator-driven plant adaptation.

## Introduction

The evolution of floral diversity in flowering plants (angiosperms) is driven by plant-pollinator interactions and their impact on the plants’ reproductive success (Cheptou 2021, Opedal 2019). The composition, abundance, and efficiency of pollinators shape the spatial distribution of floral traits within species, also known as geographic mosaics of floral morphology (Johnson 2019, Van der Niet et al. 2014). Since most plants rely on successful pollen transfer to reproduce, pollinators strongly promote diverse floral adaptations associated with traits related to mating system and attractiveness. For instance, pollinators influence pollen dispersal opportunities by facilitating mating with other plant individuals (outcrossing), while the absence of pollinators favors reproduction via self-fertilization (selfing). Therefore, pollinators play a pivotal role as selective agents in reproductive strategies (Barrett and Harder 1996). An estimated 20% of angiosperm species have switched their mating system from outcrossing to self-fertilization (Barrett 2002). The transition to a selfing mating system seems to be particularly influenced by a complex and unpredictable pollinator environment (Kalisz and Vogler 2003). The decrease or absence of effective pollinators drives selection towards selfing, along with changes in the associated phenotypic traits (Kalisz and Vogler 2003, Moeller 2006, Bodbyl Roles and Kelly 2011, Shimizu et al. 2011). For instance, in the genus *Clarkia*, two closely related species significantly differ in herkogamy (physical distance between stamen and stigma), associated with the abundance of bees in their respective pollinator communities (Moeller and Geber 2005). In *Mimulus guttatus*, the absence of pollinators led to an increase of selfing and traits promoting selfing after only five generations (Boddyl Roles and Kelly 2011). In *Brassica rapa*, successive generations of pollination by hoverflies or bumblebees resulted in divergent evolution of various plant traits (Gervasi and Schiestl 2017). However, the extent of different pollinator categories can influence the reproductive success of plants.

Due to their different functional traits (behavior, body structure, size, activities, color vision, diet specialization, etc.), different categories of pollinators (small bees, bumblebees, longue-tongue bees, hoverflies, beetles, etc.) vary in pollen transfer efficiency, affecting the adaptation of flowering plants in various ways (Harrisson et al. 2017, Hoehn et al. 2008, Dellinger 2020, Fenster et al. 2004, Albrecht et al. 2012). While bees and bumblebees are well-known for being strongly efficient pollinators, some hoverflies are considered as less-efficient pollinators (Jauker and Wolters 2008, Jauker et al. 2012). However, in an unbalanced pollinator decline – where bee and bumblebee populations decline more rapidly than hoverflies – it seems that hoverflies can play an essential role in plant pollination (Jauker and Wolters 2008, Jauker et al. 2009, Doyle et al. 2020). For instance, the hoverfly species *Episyrphus balteatus* is an important generalist pollinator for many crops and wild plant species (Jauker and Wolters 2008, Jauker et al. 2012, Doyle et al. 2020, Hodgkiss et al. 2018). While they rely on pollen as a food source as adults, they do not transfer as much pollen as some other more effective pollinators (Doyle et al. 2020). However, a reduction in the amount of pollen delivered to an outcrossing plant can lead to pollen limitation and, consequently, a decrease in reproductive success (Cheptou 2021, Gómez et al. 2010). In this context, an increase in self-pollination is predicted as an evolutionary response of flowering plants, leading to reproductive assurance (Kalisz et al. 2004). Therefore, understanding the genomic basis involved in mating system transition is paramount in predicting success and maintenance of wild and cultivated flowering plant species in the future.

Because a main driver of transition to selfing is the breakdown of self-incompatibility (SI), a substantial effort has been made to understand the underlying genetic basis, such as the molecular components of the self-incompatibility locus (SI-locus), as well as the mechanisms involved in the transition to self-compatibility (SC) (e.g., reviewed in Shimizu and Tsuchimatsu 2015). This effort was essential to understand some of the evolutionary processes involved in the frequent transition of out-crossing to selfing mating systems in angiosperms (Barrett 2002, Shimizu and Tsuchimatsu 2015). It was also of value to breeders engaged in producing homozygous lines where self-incompatibility (SI) is an undesired trait (Li et al. 2023). However, the transition to selfing is also associated with changes in a combination of phenotypic traits, known as the selfing syndrome, and only a handful of studies have unraveled their genetic basis (Tsuchimatsu and Fujii 2022). Studies in different plant species agree that the correlated evolution of floral traits observed in the selfing syndrome is likely governed by strong selection pressures (Fishman et al. 2002, Sicard and Lenhard 2011, Tsuchimatsu and Fujii 2022). Previous studies showed the involvement of a polygenic architecture underlying the selfing syndrome, including loci of both major and minor effect size (Slotte et al. 2012, Fishman et al. 2002, Goodwillie et al. 2006, Lin and Ritland 1997). In Brassicaceae, SI as well as the selfing syndrome have been widely studied and discussed, especially in *Brassica rapa*, *B. oleracea*, and *B. napus*, *Capsella* spp., *Arabidopsis thaliana*, and *A. lyrata* (Nasrallah 2017). For instance, in the genus *Capsella*, gene ontology (GO) analysis identified many genes involved in the differentiation between outcrossing and selfing in sister plant species (Woźniak et al. 2020). Altogether, the studies in *Capsella* spp. showed that the reduction of organ sizes (e.g., petal area, anther, and carpel lengths) was driven by earlier organ maturation rather than a decrease in cell number. In this case, the selfing syndrome was associated with a reduced time of cell growth (Woźniak et al. 2020). However, as far as we know, no functional analyses were performed to causally relate candidate QTLs with the regulation of organ sizes associated with the selfing syndrome.

Gervasi and Schiestl (2017) used the fast-cycling turnip (*B. rapa*) to understand the evolution of floral traits in response to pollinator efficiencies. They performed an experimental evolution study over nine consecutive generations, in three parallel replicates, with three pollination treatments: bumblebee, hoverfly, and hand pollination. Gervasi and Schiestl (2017) highlighted the importance of hoverfly pollination in the rapid evolution of the mating system to increased selfing in *B.rapa*, driven by a comparatively low efficiency in pollen transfer. Associated with the evolution of autonomous selfing, they described the evolution of selfing syndrome (reduction in most floral traits). However, we still know little about the consequences of low-efficient hoverfly pollination on the genomic architecture involved in such rapid phenotypic evolution. With a small genome size and the availability of a reference genome (Wang et al. 2011, Zhang et al. 2018), *B. rapa* is perfectly suited to study pollinator-plant interactions at the genomic level. Thus, we resurrected seeds from the first and last generations of the *B. rapa* selection experiment by Gervasi and Schiestl (2017) to study the effects of hoverflies on rapid phenotypic evolution and the underlying genomic changes. We addressed how the hoverfly-driven evolution of the transition from predominantly outcrossing to mixed mating, and the associated floral traits correlated with changes at the genomic level. Based on a genome-wide association study (GWAS), we identified candidate genes underlying floral trait variation over nine generations. We then studied the function of selected candidate genes in the regulation of floral traits by characterizing the corresponding mutants in *A. thaliana* and could confirm a role of some of them in controlling the corresponding traits. Taken together, these analyses provided novel insights into the molecular mechanisms underlying the evolution of mating systems.

## Material and methods

### Plant material

*B. rapa*, commonly known as turnip, field mustard, or Chinese cabbage, is native to the Mediterranean regions of Europe and exists in the wild and as cultivated forms (e.g., oilseed crops or vegetables like bok choy and turnip). *B. rapa* is predominantly outcrossing, which is achieved by a self-incompatibility mechanism (Hatakeyama et al. 1998). The development of a fast-cycling variety has enabled evolutionary biology research given its very short life cycle of 2 months (Tomkins and Williams 1990). An experimental evolution study was conducted on *B. rapa* with hoverfly pollination over nine generations, followed by two generations of inter-replicate crossing, which were performed to reduce and inbreeding effect, and are referred to as generation 11 (Gervasi and Schiestl 2017). For our study, we obtained *B. rapa* seeds of populations including generation 1 (before pollinator selection) and generation 11 (after hoverfly *Episyrphus balteatus* selection and inter-replicate crosses) from Gervasi and Schiestl (2017). Generation 1 was a population of highly heterozygous and outcrossing fast-cycling *B. rapa* plants comprised of 108 full-sibling seed families (Gervasi and Schiestl 2017). The hoverfly pollination treatment was divided into three replicates A, B, and C, and each replicate was independently subjected to pollinator selection for nine generations (**Figure S1**). Only pollinator-visited plants contributed to the following generation. After nine generations of selection, Gervasi and Schiestl (2017) used manual pollinations to create generation 11 by inter-replicate crosses (**Figure S1**). The inter-replicate crosses were performed on random pairs of individuals from different replicates to remove inbreeding effects. Pollinations were performed by using pollen from replicate A to pollinate replicate B, pollen from replicate B to pollinate replicate C, and pollen from replicate C to pollinate replicate A, respectively (**Figure S1**).

### Experimental design: resurrection approach

We sowed 324 seeds from generation 1 (108 full-sibling families for each of the three replicates) and 270 seeds from generation 11 (replicate A = 102 samples, replicate B = 90 samples, replicate C = 78 samples). We split the 594 plants into 11 batches of approximately 50 randomly chosen plants from both generations and their three replicates. All plants were grown under standardized conditions: first in a phytotron from day 0 to 21 days after sowing (das) (T=21°C, 24-hour light cycle, 60% relative humidity), and then in a greenhouse from 21 das to the end of the experiment (T=21°C, 16-hour light cycle, 60% relative humidity). For our study, we choose 394 samples in which both phenotypic and genomic characterization were possible, i.e., 225 samples from generation 1 (replicate A = 78 samples, replicate B = 78 samples, replicate C = 69 samples) and 169 samples from generation 11 (replicate A = 73 samples, replicate B = 45 samples, replicate C = 51 samples) for phenotypic measurement and sequencing. It should be noted that despite randomization, a batch effect was observed for all phenotypic traits (**Table S1**).

### Characterization of the mating system

We performed controlled pollination experiments to evaluate the ability of a plant to produce seeds after outcrossing (IO), selfing (SF), and autonomous selfing (AS), defined as a plant’s ability to produce seeds without manipulation. To assess the outcrossed seed production, we conducted outcrosses on flower buds one day before their presumed opening. Prior to pollination, we emasculated flower buds by removing all anthers. Then, we cross-pollinated three flowers per plant between two randomly selected plants of the same treatment. To assess the selfed seed production (*i.e.,* the ability of a plant to produce seeds through self-fertilization), we used newly opened flowers by removing anthers and using pooled pollen from the same plant to hand pollinate three flowers. To assess the autonomous selfed seed production, we labeled flower buds of the same age as the ones used for outcrossing and let flowers pollinate autonomously (without disturbance or pollinators). At harvest, we counted the number of siliques and the number of seeds produced per pollinated (labeled) flower in all mating system assays.

### Characterization of floral traits

We used the above described 394 plants of generations 1 and 11 to evaluate changes in floral traits. We collected three flowers per plant on ice for measuring floral traits at 22 das. We removed two petals and two long stamens with forceps from each flower to allow for better visualization. We taped dissected flowers along with a ruler on black paper with a double-sided tape (Scotch Double Sided Tape 12 mm x 6.3 m) and scanned them at 2,400 dots per inch using an Epson Perfection V750 PRO scanner and the Epson Scan software. We measured five floral traits using ImageJ (Version 1.53e, Waybe Rasbans and Cok NIH, 64-bit): pistil length, long stamen length, short stamen length, petal width, and distance between stigma and anther of the long stamen (herkogamy).

### Statistical analyses of phenotypic traits

We analyzed the phenotypic data in R Studio (Version 1.4.1106, PBC; R i386 4.0.4). In all tests, we considered a *p-*value below 0.05 as significant.

We evaluated the significance of phenotypic changes in response to hoverfly pollination using an ANOVA in R. We used Spearman’s (rho) ranking to establish a matrix of correlations for all floral traits and the number of seeds per flower in the controlled pollination experiments testing mating system (IO, SF, and AS) within generations 1 and 11. We used Harrell’s Miscellaneous (Hmisc) R Studio package to compute a correlation matrix (Harrell 2022) and generated heat-plots representing significant pairwise correlation coefficients for all traits. For this correlation analysis, missing values were removed (for 18 individuals from generation 1, and 11 from generation 11; a total of 365 individuals were used out of 394).

### DNA extraction and GBS sequencing

We collected and flash froze 1×1 cm^2^ of leaf tissue in liquid nitrogen from 394 plants of generations 1 and 11, and stored samples at -80°C. We performed DNA extraction and quality controls at the Genomics Diversity Center (GDC) (ETH Zurich, Switzerland). We performed tissue homogenization on a Qiagen homogenizer (frequency=300 Hz for 60 seconds, then 2 x 30 seconds), using chain tubes and two metal beads (20 grams). We used the sbeadx™ maxi plant kit by LGC group (Cat. No 41602) for DNA extraction and elution. We lysed cells, then added 1.2 µL of RNase (MN, Macherey-Nagel, Ref. 740505, 1200 U/mL, 10 mg/mL), and transferred the DNA to a 96-well KingFisher Flex plate (ThermoFischer Scientific, Waltham, USA) for extraction, using the automated magnetic-particle KingFisher Flex device at the GDC. Finally, we eluted the DNA in 80 µL of Speadex Elution buffer and assessed DNA quality using the Spark system (Tecan, Männedorf, Switzerland). We used a broad-range DNA concentration approach (2-1000 ng Quibit DNA BR buffer; Invitrogen, Waltham, USA) with following concentration standards: 0 ng (blank), 1 ng, 3 ng, 25 ng, 50 ng, and 100 ng. All samples were evaluated under 485 nm excitation and 535 nm emission wavelengths. We assessed the genomic integrity of a 1:100 dilution using a TapeStation (Agilent Technologies, Santa Clara, USA).

The 394 samples had a good DNA quality (minimum fragment size: 10 kb, conc.: ≥10-50 ng/µL). They were sequenced by LGC Genomics GmbH (Berlin, Germany) in October 2019, using Illumina NextSeq 500 V2 (Illumina, San Diego, USA) to perform GBS with the restriction enzyme *MslI*. Sequencing generated and average of 1.5 million reads for each of the 394 samples. We analyzed sequencing quality by evaluating the mapping rate to the reference sequence BrapaFPsc_277_v1 (https://phytozome-next.jgi.doe.gov/), which was at 96.38%. GBS resulted in 46,526 SNPs with a minimal read count of 8. We performed filtering, keeping SNPs present in at least 33% of the samples, with an allele frequency at or above 10% (LGC Genomics GmbH, Berlin, Germany). We removed the scaffolds and conserved the 10 chromosomes present in *B. rapa*. We finally performed a stringent quality control using VCFtools 1.12.0, keeping only the bi-allelic SNPs, discarding all SNPs with a minor allele frequency below 0.1, setting a genotype quality superior to 20, and a read depths superior to 5 (116,117). After quality control and filtering, we kept 12,254 SNPs for subsequent genomic analyses.

### Analyses of genomic changes

We performed a GWAS using the general mixed-model EMMAX (Efficient Mixed-Model Association eXpedited) to consider individual genetic relatedness (Kang et al. 2010). EMMAX includes a genetic kinship matrix implemented in PLINK 1.9 (Purcell 2007). We used five phenotypic traits (petal width, long stamen length, short stamen length, pistil length, and anther-stigma distance or herkogamy), which showed significant phenotypic evolutionary changes from generation 1 to generation 11 based on the ANOVA (**Table S1**). We evaluated SNPs of the highest association defined as the top 1% (i.e., 122 SNPs) in all further analyses.

We performed a genomic scan based on a differentiation index (using *Jost’s D* index) to uncover potential genomic signatures of selection. We calculated *Jost’s D* scores between generations 1 and 11 for 12’254 SNPs using the vcfR package (function genetic_diff) in R Studio (Version 1.4.1106, PBC; R i386 4.0.4) to quantify allelic evolution and genetic divergence (Knaus and Grünwald 2017, Jost et al. 2017). We compared the top 1% of SNPs associated with phenotypic variation (i.e., 122 SNPs with smallest *p-*values in the GWAS), with the top 10% of SNPs potentially under selection (i.e., 1225 SNPs with the highest *Jost’s D* scores) to determine whether genomic regions involved in trait variation are potentially under selection.

### Identification of candidate genes

We identified loci both involved in phenotypic trait variation (i.e., 1% SNPs (N=122 SNPs) with highest association scores in GWAS), as well as those under strongest genomic selection (i.e., 10% SNPs (N=1225 SNPs) in the tail of *Jost’s D* distribution). We also chose to include two genomic regions with significant peaks (**Figure 2A**) that were, however, not detected as under genomic selection in our analysis. In total, we analyzed 161 SNPs and retrieved the annotated genes plus surrounding 2 kb of sequence (1 kb upstream and 1 kb downstream), using the GenomicRanges package (Lawrence et al. 2013). We used the Plaza V3.0 database to identify the final 31 *B. rapa* candidate genes (https://phytozome-next.jgi.doe.gov/). Once we identified the *B. rapa* genes, we searched in *A. thaliana* databases (TAIR and NCBI) for orthologues because more mutants are available in this closely related model species. We collected data on gene function from The *Arabidopsis* Information Resource (TAIR, https://www.arabidopsis.org/), and Protein Analysis Through Evolutionary Relationships (PANTHER16.0; http://www.pantherdb.org/) classification system.

### Functional characterization of candidate genes

We assessed the involvement of the identified candidate genes in the selfing syndrome by evaluating homozygous mutants in *A. thaliana*. We used 50 homozygous (T-DNA insertion) lines obtained from the Nottingham *Arabidopsis* Stock Centre (NASC, https://arabidopsis.info/BasicForm) (**Table S2**). We grew the plants in five batches, each containing an appropriate control (i.e., wild-type Col-0). We evaluated five plants per line and nine flowers per plant for floral morphology. Flowers were collected only from the main stem and included only young flowers, defined as flowers located between the fourth and fifteenth flower counting from the bottom-up. We dissected each flower with forceps removing two petals and two long stamens to allow for better access to measuring the remaining structures. The flowers were taped on glass slides and scanned at 2400 dots per inch using an Epson Perfection V750 PRO scanner and Epson Scan software. ImageJ software (Version 1.53e, Waybe Rasbans and Cok NIH, 64-bit) was used to take measurements of floral traits from the flower scans, using the line tool at a pre-set scale (included with the scanned flowers). We performed all data analyses in the R environment and used a *t*-test to determine significant differences between wild-type control and mutant lines.

## Results

### Changes in mating system and floral traits mediated by hoverfly pollination

To understand changes in reproductive strategies over nine generations of hoverfly pollination, we characterized the ability of plants to outcross, self, and autonomously self (**Figure 1A**), as well as the variation in associated floral morphology traits. We observed significant changes in the production of outcrossed, selfed, and autonomous selfed seeds over the generations pollinated by hoverflies (**Figure 1**). For instance, we observed a drop from 17±9 to 12±7 (mean ± standard deviation) in the production of seed after outcrossing (**Table S1, Figure 1B**). In contrast, the number of seeds after self-pollination increased by 2.7-fold in selfing (**Table S1, Figure 1C**) and the ability of seed production after autonomous selfing increased from nearly zero to an average of 3.7 seeds per flower (**Table S1, Figure 1B**). These results suggested a shift from a predominantly outcrossing mating system in generation 1 to a mixed mating system (outcrossing and selfing) in the last generation mediated by hoverflies pollination (Gervasi and Schiestl 2017). This shift in mating system strategies was associated with changes in floral morphological traits. While most of the floral morphological traits were significantly positively correlated with outcrossed seed production, they were negatively correlated with the autonomous selfed seed production (**Figure S2**). Only pistil length and herkogamy were negatively correlated with the selfed seed production (**Figure S2**). Overall, we observed a significant decrease in the values of the five floral morphological traits *i.e.,* petal width, long stamen length, pistil length, and herkogamy (**Figure 2, Table S1**). For instance, we observed a reduction in pistil length from 6.25±0.85 to 5.50±0.81 cm (mean ± s.d.), and a decrease from 0.66±0.47 to 0.43±0.28 cm in herkogamy (**Figure 2, Table S1**).

**Figure 1.**
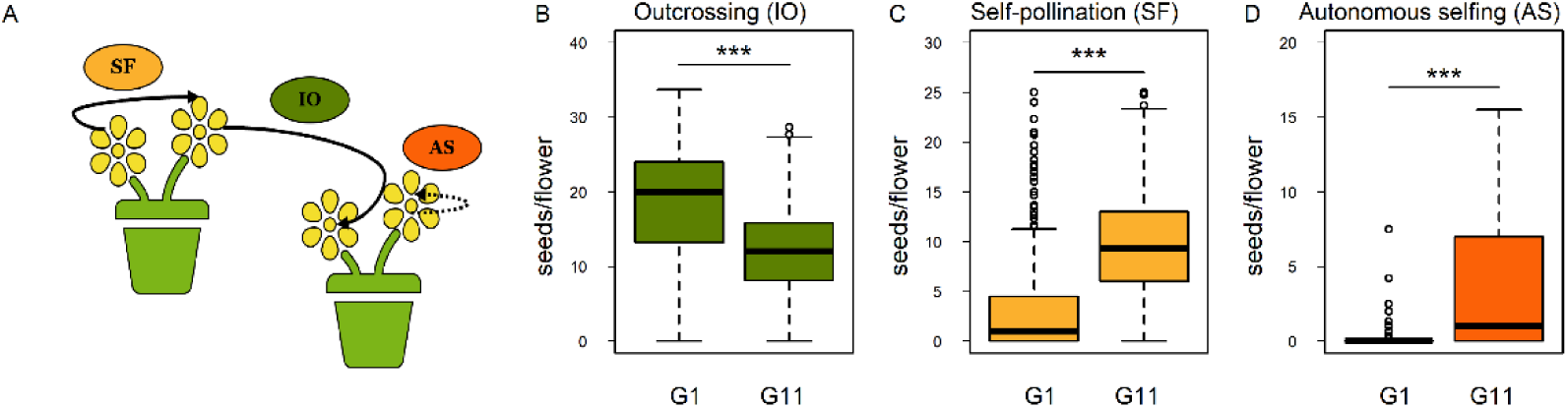
Evolution of the *B. rapa* mating system induced by hoverfly pollination. (A) Diagram of three pollination tests conducted on *B. rapa* to determine the ability to produce seeds by outcrossing (IO), self-pollination i.e., self-compatibility (geitonogamous, SF), and autonomous selfing (AS). The solid arrows indicate manual pollen transport between flowers (IO and SF), while dotted arrow indicates a potential spontaneous pollination without manual pollen transport (AS). The variation in seeds per flower produced by outcrossing (B), self-pollination (C), and autonomous selfing (D) is shown. The significant difference between generations is indicated by stars (*** for a *p-value* < 0.0001).

**Figure 2.**
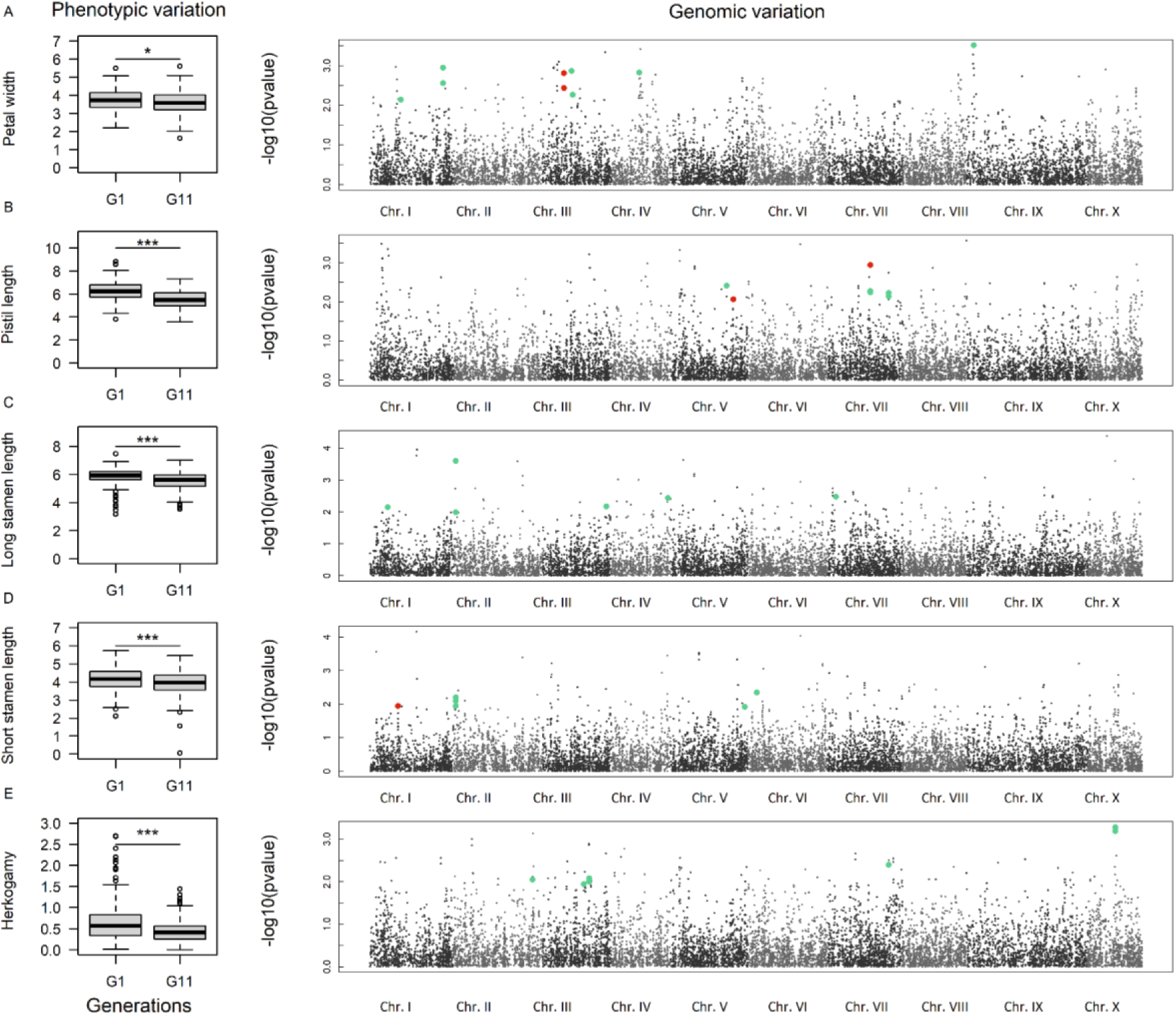
Variation in traits linked to the selfing syndrome and the associated genomic changes mediated by hoverfly pollination. The left panel shows the phenotypic variation for five floral traits: petal width (A), pistil length (B), long stamen length (C), short stamen length (D), and herkogamy (E). The *y-axis* represents the average trait in millimeters. The right panel shows the GWAs results represented by Manhattan plots. The *y-axis* shows the association score, and the *x-axis* the physical position of SNPs along the genome of *Brassica rapa.* The chromosomes are represented by gradient grey dots. Green dots represent the SNPs that belong to the identified candidate genes, and the red ones the SNPs that belong to orthologous genes functionally validated in *Arabidopsis thaliana*.

### Identification of candidate genes associated with changes in floral morphology traits

To identify genomic regions underlying the observed phenotypic changes between the first and the last generation, we performed genome-wide association (GWA) mapping on five floral morphology traits for which we observed significant changes. Based on 12’254 SNPs, we observed SNPs with high association scores for the five different traits highlighted in Manhattan plots (**Figure 2**). Several genomic regions contained SNPs that formed narrow peaks (skyscrapers); for instance (i) SNPs in the middle of chromosome IV and the beginning of chromosome IX were associated with petal width, and (ii) SNPs in the middle of chromosome X were associated with herkogamy (**Figure 2**). Using *Jost’s D* index to assess genetic differentiation, we were able to identify SNPs showing genetic differentiation between first and last generations (**Figure S3**). The maximal *Jost’s D* index observed between these two generations was *Jost’s D* = 0.8 (**Figure S3**).

We identified candidate genes in *B. rapa* based on two mutually exclusive criteria. We selected (1) either the top 1% of SNPs with the highest association scores in GWAS and overlapping with the top 10% of SNPs with the highest *Jost’s D* scores, or (2) SNPs within a narrow skyscraper in GWAS but excluded from the top 10% of SNPs with highest *Jost’s D* index. Overall, we identified 31 candidate genes, for which we could identify orthologous genes in *A. thaliana* and their associated biological function (**Table S2, Table S3**).

### Functional analysis of orthologous genes in A. thaliana

Based on the identified candidate genes in *B. rapa*, we performed a functional analysis using mutants of orthologues in *A. thaliana*. We analyzed a total of 50 publicly available homozygous T-DNA insertion mutants of these orthologues (**Table S2**). We found significant differences between the wild type and four mutants involved in phenotypic changes for petal width (one mutant; N684414 associated with *AT2G31010*), pistil length (two mutants; N662707 and N656817 associated with *AT3G14980* and *AT2G36620* respectively), and short stamen length (one mutant; N678039 associated with *AT4G17080*) (**Figure 3**, **Table 1**). We did not observe any significant phenotypic differences between wild-type and mutant plants for candidate genes involved in herkogamy or long stamen length (**Figure S4**). In detail, we identified one mutant affecting a gene involved in petal size (**Table S3, Figure 4A**). This gene encodes a protein kinase annotated as involved in the response to organic cyclic compounds. We identified two mutants whose pistil lengths significantly differed from the wild type (**Table S3, Figure 4B**). The first mutant disrupts the *INCREASED DNA METHYLATION1*/*REPRESSOR OF SILENCING4* (*IDM1*/*ROS4*) gene, which is involved in gene silencing via the RNA-directed DNA-methylation pathway (**Table S3**). The second affects the ribosomal protein L24 involved in protein synthesis (cytoplasmic translation) (**Table S3**). Finally, differences in short stamen length were found in homozygous *Arabidopsis thaliana reduction in growth and productivity* (*Atrgp*) mutants, affecting a SET-domain containing H3K4-specific histone methyltransferase involved in defense responses (**Table S3, Figure 3C**).

**Figure 3.**
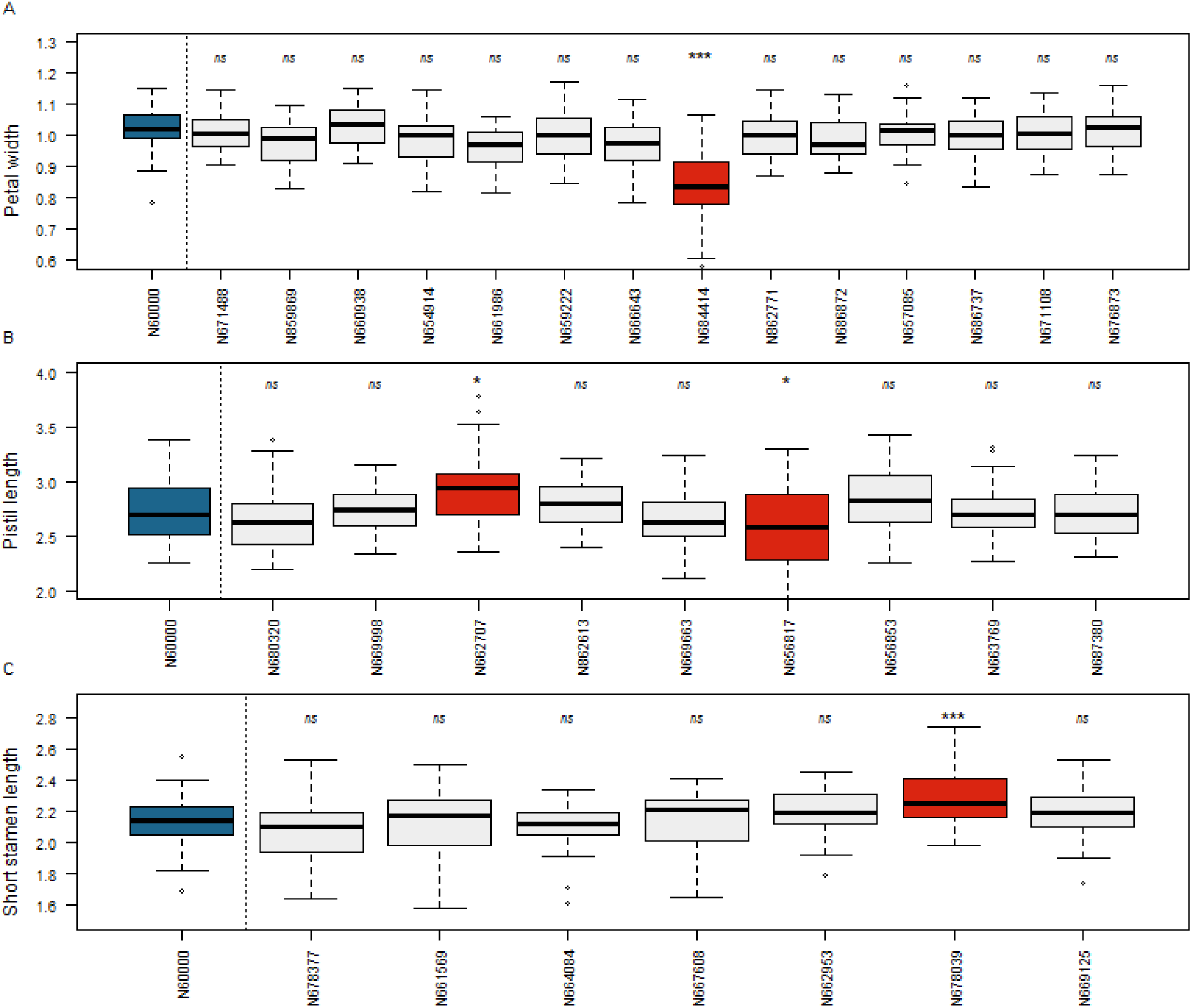
Functional analysis of *Arabidopsis thaliana* mutants. Floral phenotype variation of the mutants compared to the wild-type (Col-0) for petal width (A), pistil length (B) and short stamen length (C). Herkogamy and long stamen length are shown in Figure S4. The *y-axis* represents the phenotypic trait variation in millimeters, and the *x-axis* shows the wild-type and the different mutant lines. The wild-type (Col-0, N60000) is represented by a blue whisker box, while mutant lines with a significant difference to the wild type are represented by red whisker boxes. The significance of the pairwise t-test between wild-type and mutant lines is indicated above each whisker box: *ns* for non-significant, * for *pvalue* < 0.05, ** for *pvalue* < 0.01, *** for *pvalue* < 0.001.

**Table 1.** Among 31 candidate genes identified in *Brassica rapa*, four orthologous candidate genes in *Arabidopsis thaliana* were functionally validated. The GO terms, Biological Process, were retrieved from the TAIR database. Only genes with significant statistical difference in floral organ sizes in the corresponding mutant are shown in the table. For a full list of results see Table S3.

| Trait in association | *A. thaliana* identifier | Gene name | Gene function (GO biological term) |
| --- | --- | --- | --- |
| Petal width | AT2G31010 | Protein kinase superfamily protein | phosphorylation, response to organic cyclic compound; protein phosphorylation |
| Pistil length | AT3G14980 | REPRESSOR OF SILENCING 4 (ROS4) - Acyl-CoA N-acyltransferase with RING/FYVE/PHD-type zinc finger protein | gene silencing by RNA-directed DNA methylation, histone H3-K14 acetylation, histone H3-K18 acetylation, histone H3-K23 acetylation, regulation of DNA methylation |
| Pistil length | AT2G36620 | Ribosomal protein L24 (RPL24A) | response to inorganic substance; cytoplasmic translation |
| Short stamen length | AT4G17080 | Histone H3 K4-specific methyltransferase SET7/9 family protein | defense response to other organism |

## Discussion

We investigated the impact of low-efficient pollinators on rapid plant evolution by identifying 31 candidate genes underlying changes related to the selfing syndrome *Brassica rapa* after nine generations of hoverfly pollination. Furthermore, we functionally validated the role of four candidate genes in selfing syndrome traits in *B. rapa* by studying the corresponding mutants disrupting their *A. thaliana* homologues.

### Evolution of phenotypic strategies associated with a mixed mating system

We confirmed the finding of Gervasi and Schiestl (2017) that nine generations of hoverfly pollination led to the transition of a primarily outcrossing to a mixed mating system (i.e., both a decrease of outcrossed seed production, an increase in self-compatibility, and an increase in autonomous selfing) in *B. rapa*. These findings are aligned with prior theorical approaches and observations where the shift from self-incompatible plant species to a mixed mating system was documented under an unstable pollinator environment (Kalisz et al 2004, Cheptou and Massol 2009, Eckert et al. 2010, Thomann et al. 2013). Consistent with the prevalence of mixed mating systems in flowering plants (Goodwillie et al. 2005, Whitehead et al. 2018), mixed mating systems can provide the dual advantage of promoting outcrossing and selfing in the short-term. While the rate of outcrossing effectively maintains genetic diversity within populations and reduces inbreeding, the ability to self-fertilize ensures reproductive assurance. However, a high selfing rate could have a negative effect in the long-term maintenance of populations due to genomic constraints (Cheptou 2019). Therefore, this mixed mating system represents an outcrossing-selfing paradigm with a critical balance between genetic variation and reproductive assurance (Barrett 2003) and appears to be highly variable at the intraspecific level (Whitehead et al. 2018). In the current context of insect pollinator decline, an increase in mixed mating populations with an increase in autonomous selfing is expected (Kalisz and Vogler 2003, Shivanna 2015). The complex interaction between pollinators and the dynamics of changing mating systems is reflected in subsequent phenotypic adaptations of the floral morphology.

Our study unveiled a shift to a mixed mating system associated with changes in floral phenotypic strategies. These changes were characterised by traits related both to resource allocation reduction in outcrossing, such as reduced petal width, and traits reducing the efficiency of the SI mechanism, such as a decrease in the length of reproductive organs and herkogamy. These changes in response to hoverfly pollination are characteristic of the selfing syndrome that has been described in many species (e.g., in the genera *Arenaria, Capsella, Clarkia, Collinsia, Eichhornia, Leptosiphon, Mimulus, Ipomea,* and *Solanum*) and is common during a shift of the mating system from predominantly outcrossing to selfing (Sicard et al. 2016, Fishman et al. 2002, Fishman and Willis 2008, Sicard and Lenhard 2011, Goodwillie et al. 2006, Lin and Ritland 1997, Georgiady et al. 2002, Vallejo-Marín and Barrett 2009, Ducan and Rausher 2013, Tsuchimatsu and Fujii 2022). Furthermore, the rapid evolution of the selfing syndrome could be triggered by the decline of pollinators in natural populations, as recently observed in *Viola arvensis* over a 20- year period (Cheptou et al. 2022). To predict the evolution of mating systems in natural populations of flowering plants in the context of pollinator change, it is essential to better understand the genetic basis involved in the traits associated with the selfing syndrome.

### Multi-locus basis of selfing syndrome evolution in hoverfly-pollinated plants

Our study revealed several genes involved in the selfing syndrome associated with the adaptive response of plants to hoverflies. In total, we identified 31 candidate genes associated with the reduction of floral organ size and herkogamy. These candidate genes were mostly involved in cell-to-cell signaling, transport, and the development of plant organs (Kaur et al. 2006, Chua et al. 2005, Wang et al. 2012, Tronconi et al. 2008, Liu et al. 2013). Despite our relatively limited set of markers for the GWA mapping, we validated the functional role of four *A. thaliana* homologues involved in regulating the size of floral organs relevant to the selfing syndrome. The first functionally validated gene was involved in petal size and encoded a protein kinase, indicating a regulatory role (**Table 1**, **Figure 2A**). In plant-biotic interactions, protein kinases have important roles in the perception of insects and signal transmission in plant-herbivore interactions (Romero-Hernandez and Martinez 2022). While the role of protein kinases is not well documented in plant-pollinator interactions, studies have demonstrated their roles in petal shape (a trait in plant attractiveness to pollinators) in *Brassica napus* (*e.g.*, cyclin-dependent kinase inhibitor ICK1; Zhou et al 2002) or in *A. thaliana* (e.g., ERECTA receptor kinase; Abraham et al. 2013). The second and third candidate genes validated were involved in the variation of pistil length. We validated a ribosomal protein (L24A) gene to be associated with the variation in pistil length. Ribosomal proteins have an important role in the growth and development of plants. For instance, a high expression of this gene in flowers during osmotic stress has been found (Park et al. 2017). In addition, we identified a gene encoding the protein INCREASED DNA METHYLATION1 (IDM1)/REPRESSOR OF SILENCING4 (ROS4), involved in the regulation of active DNA demethylation (**Table 1**, **Figure 2B**, Li et al. 2012). Together with other components, IDM1/ROS4 forms a complex to recruit the DNA demethylation enzyme ROS1 and is involved in a broad range of developmental processes and responses to biotic and abiotic stresses (Li et al. 2018, Liu and Lang 2020). Finally, we validated the function of the Histone H3 K4-specific methyltransferase SET7/9 in controlling the length of the short stamen. Given that the short stamens were longer in the corresponding mutant but became shorter during hoverfly pollination over nine generations, it is likely that SET7/9 is a repressor of short stamen length, and its expression was higher in the last as compared to the first generation. Interestingly, the previously described ROS4/IDM1 protein, involved in pistil length, binds to the unmethylated histone H3K4 involved in short stamen length (Li et al. 2012, Qian et al. 2012). It is fascinating to observe that the traits involved in herkogamy (pistil and stamen length) can be controlled by genes encoding proteins involved in the same pathway.

## Conclusions

Reproductive success of flowering plants depends on the efficiency of their mating systems, determined, among other factors, by their pollinator communities. Our study showed that hoverfly pollination can lead to genomic changes associated with a shift in the mating system from predominantly outcrossing to mixed mating. To further our understanding of the molecular mechanisms involved, future work including whole-genome sequencing and genome scans on a larger scale could significantly expand our current list of candidate genes. In addition, further functional analyses in *Brassica rapa* could be performed on the already identified candidate genes to unravel the mechanism(s) underlying mating system shifts more thoroughly.

## Acknowledgements

We thank Kentaro Shimizu and his lab members (University of Zurich), Karl Schmid (University of Hohenheim), and John Pannell (University of Lausanne) for discussions and advice. We are grateful to the GDC and LGC teams for assistance with DNA extraction and sequencing, NASC for *A. thaliana* seeds, the gardeners Markus Meierhofer and Rayko Jonas (University of Zurich) for their outstanding support, the laboratory and greenhouse management teams in the Grossniklaus group, and the coordinators of the University Research Priority Project “Evolution in Action”, Yvonne Steinbach, Annegret Lessauer, Cornelia Roder, and Mira Portmann, for all their support in managing this project. Finally, we thank Andreas Kofler for aid in seed and data processing, and Laura Piñeiro for comments on the manuscript. This work was supported by the University of Zurich, a grant for a PhD project from the University Research Priority Program “Evolution in Action” to F.S. and U.G, a grant from the Julius Klaus Stiftung to X.K. and U.G., and core funding of the University of Zurich to U.G.

## Competing interests

The authors declare that they have no competing interests.

## Authors’ contributions

Conceived the project: F.S. and U.G. Raised funding: F.S., U.G., and X.K. Supervised the project: F.S., L.F., and U.G. Designed experiments: U.G., F.S., X.K., and L.F. Performed experiments: X.K. Analyzed data: X.K. and L.F. Wrote the paper: X.K and L.F., with contributions from U.G. and F.S. Coordinated the project: X.K. and L.F.

## Data Accessibility

The data and materials used in this study are available by request from the corresponding author.

## Supporting Information

**Figure S1.** Experimental evolution design.

**Figure S2.** Spearman correlations between mating system and floral traits.

**Figure S3.** Genome scan of fixation index (*Jost’s D*) between first and last generation.

**Figure S4.** Functional analysis of *Arbidopsis thaliana* mutants.

**Table S1.** Variation of floral traits after nine generations of pollination by hoverflies.

**Table S2.** 31 *Brassica rapa* candidate genes, their *Arabidopsis thaliana* orthologs, and the ID of the mutant lines from the Nottingham Arabidopsis Stock Centre (NASC).

**Table *S3****. Arabidopsis thaliana* orthologs of the candidate genes and their functional classification derived from the Plaza and TAIR databases.

## Notes

### Competing Interest Statement

The authors have declared no competing interest.

