## Supplementary Information for "Uncovering genes involved in pollinator-driven mating system shifts and selfing syndrome evolution in *Brassica rapa*"

**New Phytologist**
Full paper

### Supplementary Information

**Title.** Uncovering genes involved in pollinator-driven mating system shifts and selfing syndrome evolution in Brassica rapa

**Authors:** Xeniya V. Kofler, Ueli Grossniklaus, Florian P. Schiestl, Léa Frachon

**Figure S1. Experimental evolution design.** We used hoverfly-selected populations of *Brassica rapa* generated in an experimental evolution study previously published by Gervasi and Schiestl (2017). In their study, 108 full-sibling fast-cycling *B. rapa* families were used. The hoverfly treatment was subdivided into three replicates (A, B, and C) with 36 individuals each for nine consecutive generations. Generations 10 and 11 were created by inter-replicate crosses. In the current study, we resurrected seeds from generations 1 and 11 (3 replicates), phenotyped then, and sequenced all 394 individuals to perform a genome-wide association study (GWAS).

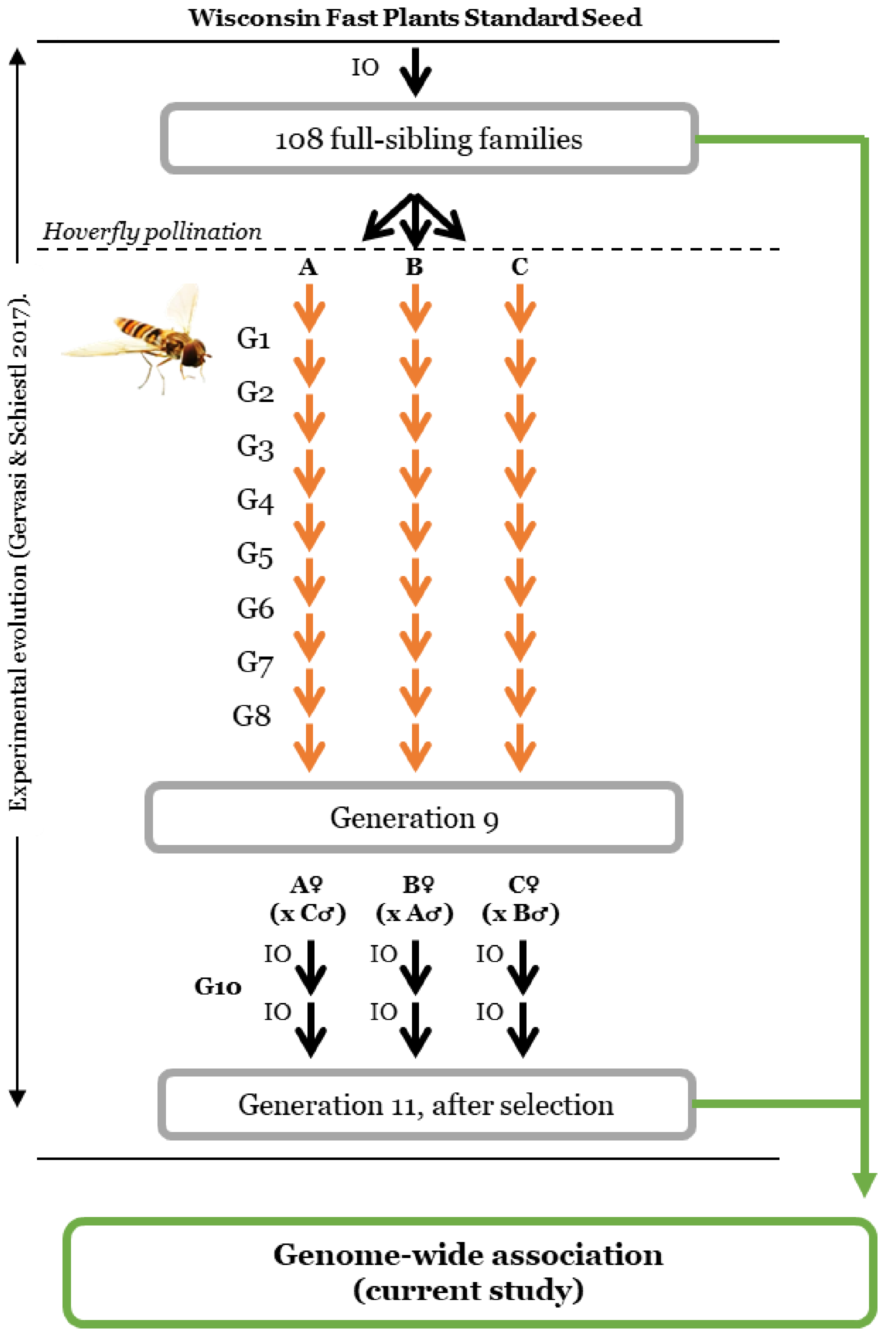

**Figure S2. Spearman correlations between mating system and floral traits.** Spearman's rho ranking matrix among mating system assessment values (seeds per flower via outcrossing, self-pollination, and autonomous selfing) and floral traits. Both generations are considered. The significant correlation values (-1 to 1) are indicated by a color gradient shown in the lower part of the plots (negative correlations are in red; positive correlations are in blue) as well as the size of circles. Non-significant correlations are indicated by the absence of a circle.

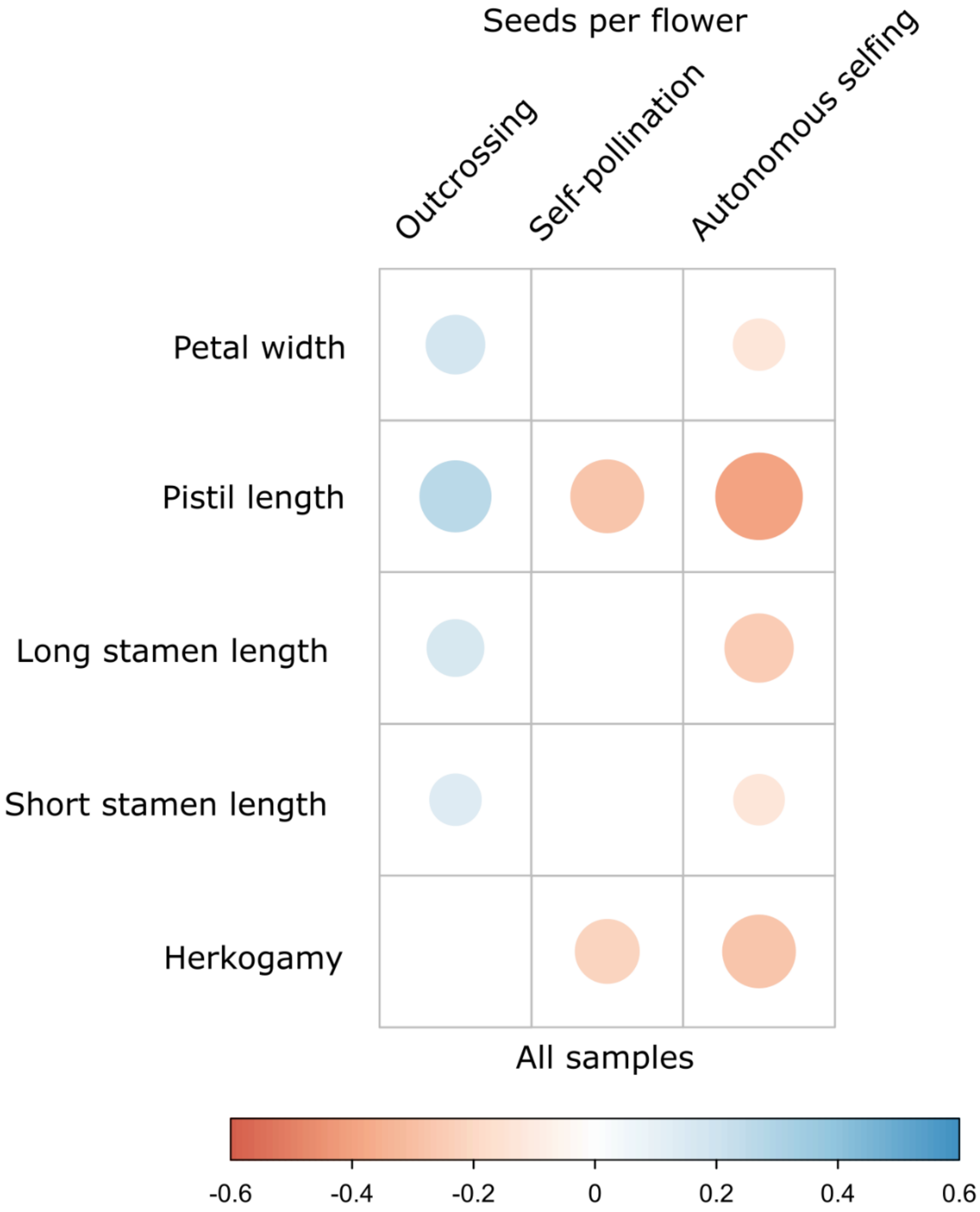

**Figure S3. Genome scan of fixation index (*Jost's D*) between first and last generation.** The Manhattan plot represents *Jost's D* scores (*y-axis*) over the ten chromosomes of *Brassica rapa* (*x-axis*). The blue dots indicate the top 10% of SNPs with the highest scores.

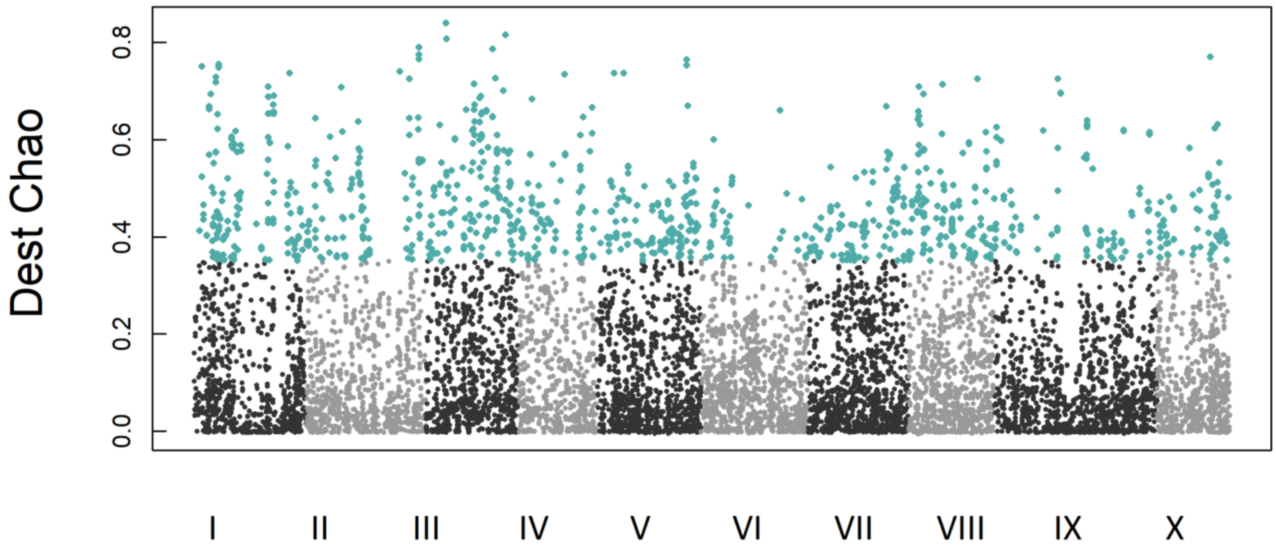

**Figure S4. Functional analysis of *Arbidopsis thaliana* mutants.** The floral phenotype of the mutants compared to the wild-type (Col-0, N60000) for herkogamy (A) and long stamen length (B). The *y-axis* represents the phenotypic trait variation in millimeters, and the *x-axis* represents the wild-type and the different mutant lines. The wild-type is represented with a blue whisker box. The significance of the pairwise t-test between wild type and mutant lines is indicated above each whisker box: *ns* for non-significant, * for *pvalue* < 0.05, ** for *pvalue* < 0.01, *** for *pvalue* < 0.001.

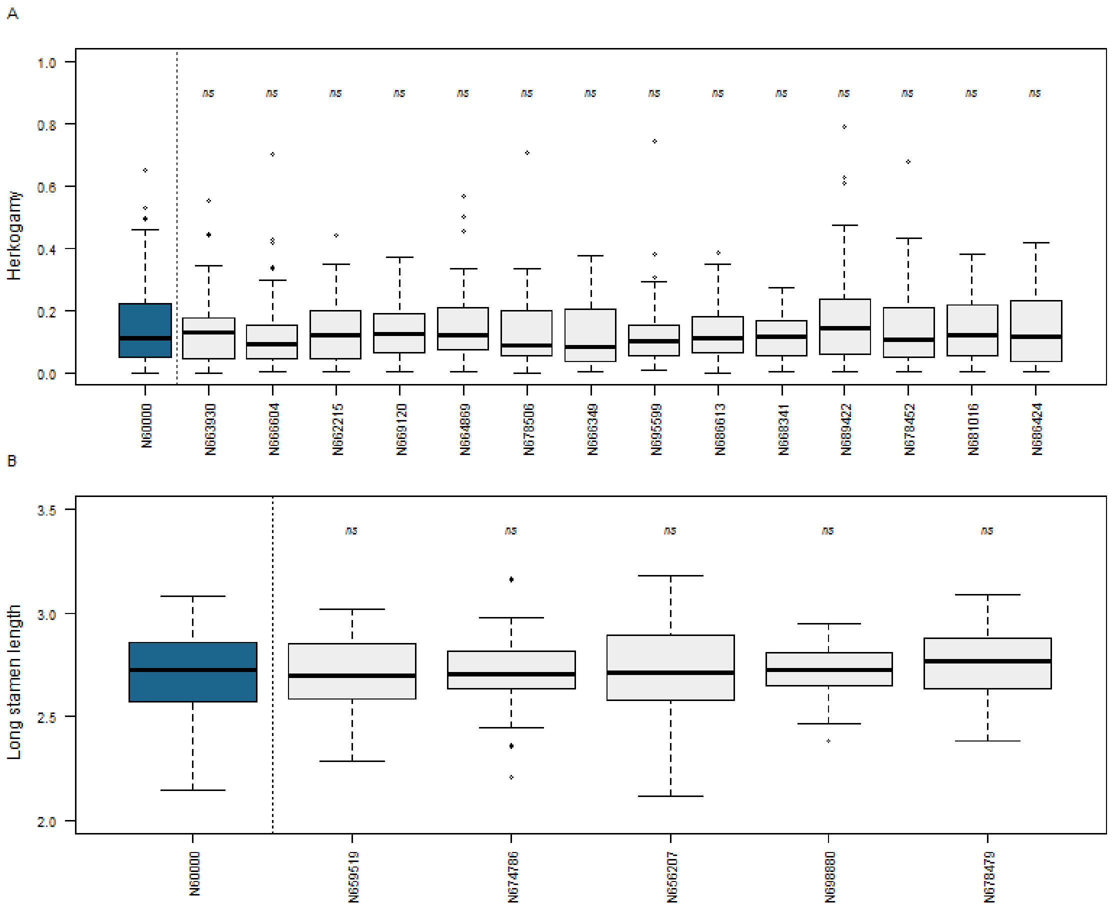

**Table S1. Variation of floral traits after nine generations of pollination by hoverflies.** The mean ± standard deviation (sd) of the measured traits in generations 1 and 11 are indicated. The effect 'generation' and 'batch' were tested using an ANOVA test (aov), and the significances are indicated as follows: *p*-values, *p*; * *P* < 0.05, ** *P* < 0.01, *** *P* < 0.001).

| | | Generation 1 | | Generation 11 | Generation effect | | Batch effect | |
|---|---|---|---|---|---|---|---|---|
| | | mean trait ± sd | | mean trait ± sd | *F value* | *P* | *F value* | *P* |
| Reproductive traits | | | | | | | | |
| **seeds/flower** | **outcrossing** | 17.43 ± 8.89 | ↘ | 12.20 ± 6.91 | 38.63 | *** | 2.73 | *** |
| | **selfing** | 3.62 ± 5.71 | ↗ | 9.87 ± 5.93 | 106.6 | *** | 3.75 | *** |
| | **autonomous** | 0.12 ± 0.66 | ↗ | 3.71 ± 4.42 | 131.6 | *** | 3.86 | *** |
| Flower morphological traits | | | | | | | | |
| **Petal width** | | 3.73 ± 0.60 | ↘ | 3.57 ± 0.67 | 5.88 | * | 3.59 | *** |
| **Pistil length** | | 6.25 ± 0.85 | ↘ | 5.50 ± 0.81 | 71.11 | *** | 7.07 | *** |
| **Long stamen length** | | 5.83 ± 0.66 | ↘ | 5.51 ± 0.65 | 21.83 | *** | 7.31 | *** |
| **Short stamen length** | | 4.16 ± 0.62 | ↘ | 3.91 ± 0.72 | 12.73 | *** | 4.1 | *** |
| **Herkogamy** | | 0.66 ± 0.47 | ↘ | 0.43 ± 0.28 | 30.08 | *** | 3.06 | *** |

**Table S2. 31 *Brassica rapa* candidate genes, their *Arabidopsis thaliana* orthologs, and the ID of the mutant lines from the Nottingham Arabidopsis Stock Centre (NASC).** The asterisks on associated traits indicate that the SNP is in the 1% with the highest scores in the GWAS, but not in the 10% SNPs with highest *Jost's D* index. The NASC IDs are the used mutant lines, and an (NA) follows the ones that were not available. The bold lines are the SNPs with a functionally validated role in the corresponding trait.

| Associated trait | SNP ID | Identifier *B. rap* | *A. thaliana* ortholog | NASC ID |
|---|---|---|---|---|
| Petal width | 1_11510427 | Brara.A01973 | AT1G51990 | N671488 |
| | | | AT1G51990 | N859869 |
| Petal width | 3_10472925 | Brara.C02099 | AT2G40980 | N660938 |
| | | | AT2G40980 | N654914 |
| Petal width | 3_10910305 | Brara.C02174 | AT2G42890 | N661986 |
| | | | AT2G42890 | N659222 |
| | | | AT2G42890 | N666643 |
| Petal width | 1_26948260 | Brara.A03224 | AT3G13940 | NA |
| Petal width | 1_26948260 | Brara.A03225 | AT3G13920 | NA |
| **Petal width** | **3_7697404** | **Brara.C01582** | **AT2G31010** | **N684414** |
| | | | AT2G31010 | N862771 |
| | | | AT2G31010 | N686872 |
| Petal width* | 9_3005754 | Brara.I00536 | AT5G25610 | N864500 (NA) |
| | | | AT5G25610 | N657085 |
| | | | AT5G25610 | N686737 |
| Petal width* | 4_10212651 | Brara.D01100 | AT5G40840 | N671108 |
| | | | AT5G40840 | N865412 (NA) |
| | | | AT5G40840 | N676873 |
| Pistil length | 5_20653707 | Brara.E02263 | AT3G43960 | N680320 |
| Pistil length | 5_23163589 | Brara.E02639 | AT1G53290 | N669998 |
| | | | AT1G53290 | N664343 (NA) |
| **Pistil length** | **5_23163589** | **Brara.E02638** | **AT3G14980** | **N662707** |
| Pistil length | 7_22322180 | Brara.G02762 | AT1G68100 | N862613 |
| | | | AT1G68100 | N669663 |
| **Pistil length** | **7_15627113** | **Brara.G01568** | **AT2G36620** | **N656817** |
| | | | AT2G36620 | N656853 |
| | | | AT3G53020 | N663769 |
| | | | AT3G53020 | N661290 (NA) |
| Pistil length* | 7_15627113 | Brara.G01569 | AT3G53110 | NA |
| Pistil length | 7_15625468 | Brara.G01567 | AT3G52990 | N687380 |
| Long stamen length | 3_23215892 | Brara.C04410 | AT3G52280 | N659519 |
| Long stamen length | 7_3006589 | Brara.G00312 | AT2G16780 | N674786 |
| Long stamen length | 3_23215892 | Brara.C04409 | AT4G38160 | N656207 |
| Long stamen length | 4_20683960 | Brara.D02701 | AT2G44600 | N698880 |
| Long stamen length | 1_6625930 | Brara.A01246 | AT4G22260 | N678479 |
| Short stamen length | 5_27296000 | Brara.E03427 | AT3G04740 | N678377 |
| Short stamen length | 6_3311183 | Brara.F00556 | AT1G08800 | N661569 |
| | | | AT1G08800 | N664084 |
| | | | AT1G08800 | N667608 |
| **Short stamen length** | **1_10431879** | **Brara.A01835** | AT4G17080 | N662953 |
| | | | **AT4G17080** | **N678039** |
| Short & long stamen length | 2_622890 | Brara.B00123 | AT5G03960 | N669125 |
| | | | AT5G03960 | N863398 (NA) |
| Short & long stamen length | 2_622890 | Brara.B00122 | NA | NA |
| Herkogamy | 2_28844740 | Brara.B03444 | AT5G48560 | N663930 |
| | | | AT5G48560 | N666604 |
| Herkogamy | 3_14912861 | Brara.C02902 | AT4G00570 | N662215 |
| | | | AT4G00570 | N860677 (NA) |
| | | | AT4G00570 | N669120 |
| Herkogamy | 3_16903670 | Brara.C03349 | AT4G31680 | N680700 (NA) |
| | | | AT4G31680 | N664869 |
| | | | AT4G31680 | N678506 |
| | | | AT4G31680 | N666349 |
| | | | AT5G32460 | N695599 |
| | | | AT5G32460 | N686613 |
| Herkogamy | 7_22282246 | Brara.G02749 | AT1G67930 | N668341 |
| | | | AT1G67930 | N689422 |
| Herkogamy* | 10_10005046 | Brara.J01029 | AT5G55920 | N678452 |
| | | | AT5G55920 | N681016 |
| | | | AT5G55920 | N686424 |
| Herkogamy* | 10_10005046 | Brara.J01030 | AT5G55940 | N2105368 |

**Table** ***S3. Arabidopsis thaliana*** **orthologs of the candidate genes and their functional classification derived from the Plaza and TAIR databases.** Bold text: genes with a demonstrated role in the corresponding trait.

| Associated trait | *A. thaliana* ortholog | *A. thaliana* full name, symbol, description (Plaza) | GO biological term (TAIR) |
|---|---|---|---|
| Petal width | AT1G51990 | o-methyltransferase family protein | - |
| Petal width | AT2G40980 | protein kinase superfamily protein | carbohydrate derivative metabolic process, cell wall organization or biogenesis, seed development |
| Petal width | AT2G42890 | MEI2-like 2 (ML2) | positive regulation of growth, positive regulation of meiotic nuclear division |
| Petal width | AT3G13940 | DNA binding / DNA-directed RNA polymerase | RNA polymerase I preinitiation complex assembly, transcription elongation from RNA polymerase I promoter |
| Petal width | AT3G13920 | eukaryotic translation initiation factor 4A1 (EIF4A1) | cytoplasmic translational initiation |
| **Petal width** | **AT2G31010** | protein kinase superfamily protein | phosphorylation, response to organic cyclic compound; protein phosphorylation |
| Petal width* | AT5G25610 | RESPONSIVE TO DESSICATION 22 (RD22) | response to abscisic acid, response to desiccation, response to salt stress |
| Petal width* | AT5G40840 | Rad21/Rec8-like family protein (SYN2) | double-strand break repair, mitotic cell cycle; replication-born double-strand break repair via sister chromatid exchange |
| Pistil length | AT3G43960 | cysteine proteinases superfamily protein | root hair elongation |
| Pistil length | AT1G53290 | galactosyltransferase family protein | cell differentiation, cell wall biogenesis, cell wall organization or biogenesis, plant epidermis development, plant-type cell wall organization or biogenesis, positive regulation of cellular biosynthetic process, positive regulation of nucleobase-containing compound metabolic process, tissue development, xylan metabolic process; protein glycosylation |
| **Pistil length** | **AT3G14980** | REPRESSOR OF SILENCING 4 (ROS4) - Acyl-CoA N-acyltransferase with RING/FYVE/PHD-type zinc finger protein | gene silencing by RNA-directed DNA methylation, histone H3-K14 acetylation, histone H3-K18 acetylation, histone H3-K23 acetylation, regulation of DNA methylation |
| Pistil length | AT1G68100 | IAA-ALANINE RESISTANT 1 (IAR1) - ZIP metal ion transporter family | zinc ion transmembrane transport |
| **Pistil length** | **AT2G36620** | ribosomal protein L24 (RPL24A) | response to inorganic substance; cytoplasmic translation |
| Pistil length* | AT3G53110 | LOW EXPRESSION OF OSMOTICALLY RESPONSIVE GENES 4 (LOS4) - p-loop containing nucleoside triphosphate hydrolases superfamily protein | poly(A)+ mRNA export from nucleus, response to abscisic acid, response to cold, response to heat; poly(A)+ mRNA export from nucleus |
| Pistil length | AT3G52990 | Pyruvate kinase family protein | - |
| Long stamen length | AT3G52280 | general transcription factor group E6 (GTE6) | chromatin remodeling; regulation of transcription, DNA-templated |
| Long stamen length | AT2G16780 | MULTICOPY SUPPRESSOR OF IRA1 2 (MSI2) - transducin family protein / WD-40 repeat family protein | - |
| Long stamen length | AT4G38160 | pigment defective 191 (pde191) - mitochondrial transcription termination factor family protein | chloroplast organization, tRNA processing |
| Long stamen length | AT2G44600 | hypothetical protein | defense response to bacterium, immune system process, regulation of defense response, response to fungus, response to nitrogen compound, response to osmotic stress, signal transduction |
| Long stamen length | AT4G22260 | IMMUTANS (IM) -alternative oxidase family protein | carotenoid biosynthetic process, chloroplast organization, plastid organization, response to high light intensity, response to temperature stimulus |
| Short stamen length | AT3G04740 | STRUWWELPETER (SWP) - RNA polymerase II transcription mediator | abscisic acid-activated signaling pathway, cold acclimation, positive regulation of cell population proliferation, positive regulation of transcription, DNA-templated, systemic acquired resistance; positive regulation of transcription by RNA polymerase II |
| Short stamen length | AT1G08800 | MyoB1,myosin binding protein 1 | response to abscisic acid |
| **Short stamen length** | AT4G17080 | Histone H3 K4-specific methyltransferase SET7/9 family protein | defense response to other organism |
| Short & long stamen length | AT5G03960 | IQ-domain 12 (IQD12) | - |
| Short & long stamen length | NA | NA | NA |
| Herkogamy | AT5G48560 | basic helix-loop-helix (bHLH) DNA-binding superfamily protein (CIB2, CRY2-interacting bHLH 2) | regulation of transcription, DNA-templated, response to blue light |
| Herkogamy | AT4G00570 | NAD-dependent malic enzyme 2 (NAD-ME2) | malate metabolic process |
| Herkogamy | AT4G31680 | Transcriptional factor B3 family protein | - |
| Herkogamy | AT1G67930 | Golgi transport complex protein-like protein | intra-Golgi vesicle-mediated transport |
| Herkogamy* | AT5G55920 | OLIGOCELLULA 2 (OLI2) - S-adenosyl-L-methionine-dependent methyltransferases superfamily protein | cell division, leaf morphogenesis, root morphogenesis; maturation of LSU-rRNA, rRNA base methylation |
| Herkogamy* | AT5G55940 | embryo defective 2731 (emb2731) - Uncharacterized protein family | embryo development ending in seed dormancy |
